# Effects of dietary replacement of broiler litter with *Melia azedarach* foliage on productive performance, fatty acid composition and health-related fatty acid indices in beef fats from Nguni x Brahman steers fed sugarcane tops based diets

**DOI:** 10.1101/573667

**Authors:** D.M.N. Mthiyane, B.J. Dlamini, A. Hugo, I.V. Nsahlai

**Affiliations:** Department of Animal Science, Faculty of Agriculture, University of Swaziland, Luyengo, M205, Swaziland; Department of Microbial, Biochemical and Food Biotechnology, Faculty of Natural and Agricultural Sciences, University of the Free State, Bloemfontein, 9300, South Africa; Discipline of Animal and Poultry Sciences, School of Agricultural, Earth and Environmental Sciences, University of KwaZulu-Natal, Pietermaritzburg, Scottsville, 3209, South Africa

**Keywords:** *Melia azedarach* leaf meal, nutritional composition, performance, fatty acid composition, disease indices, beef cattle

## Abstract

The study investigated the nutritional value of *M. azedarach* Linn. (umsilinga; *Meliaceae* family) leaf meal (MALM) as an alternative protein source for beef cattle. In a completely randomised design (CRD), 18 Nguni x Brahman 18–20 months old steers were randomly offered 3 iso-energetic and iso-nitrogenous dietary treatments with, respectively, 0% (Control), 15% and 30% MALM replacing broiler litter, each with 6 replicate animals, for 90 days. Feed intake (FI), water intake (WI), slaughter weight, body weight gain (BWG), feed conversion efficiency (FCE) and kidney fat depot fatty acid (FA) composition as well as health-related FA totals, ratios and other indices were measured. Results showed MALM contained rather high CP (290.0 g/kg DM), CF (170 g/kg DM), EE (78.1 g/kg DM) and ash (77.5 g/kg DM) contents. Also, dietary supplementation with MALM increased BWG and FCE (P < 0.01) but had no effect on FI, WI and slaughter weight of steers (P > 0.05). Also, it increased kidney fat margaric acid (P < 0.05) whilst it decreased arachidic acid (P < 0.01) content. There was no effect of diet on the content of all other saturated fatty acids (SFAs) (P > 0.05) in kidney fat. Further, dietary MALM supplementation increased kidney fat α-linolenic (P < 0.001) and conjugated linoleic acid (CLA) (P = 0.01) but had no effect on the content of all other unsaturated fatty acid (UFA) components (P > 0.05). Furthermore, it increased kidney fat total polyunsaturated fatty acids (PUFAs) (P < 0.01), total omega-3 (*n*-3) PUFAs (P < 0.001) and the CLA/vaccenic acid index but decreased the omega-6/omega-3 (*n*-6 PUFA/*n*-3 PUFA) ratio (P = 0.001). Otherwise, dietary MALM supplementation had no effect on all other FA totals, ratios and health-related indices (P > 0.05). In conclusion, dietary MALM supplementation improved productive performance of beef steers and enhanced their desirable meat fat FA profiles.

**Implications:** Broiler litter is widely used as an alternative cheap protein and mineral rich feedstuffs for supplementing poor quality forage based diets for ruminants in Southern Africa and elsewhere. However, its use is limited by the presence of human pathogens, pesticides, drug residues and heavy metals, which pose health hazards to livestock and human consumers. Our study demonstrated MALM as a better and safer alternative source of protein, the use of which in beef cattle diets would improve animal productivity and the desirable FA profile of meat which would potentially enhance the health status of consumers. By improving beef productivity, dietary MALM supplementation thus promises to enhance food and nutrition security and to contribute to poverty alleviation in Eswatini, Africa and beyond.

## Introduction

The world human population has almost doubled over the past three decades with corresponding increase in world food production (Tian *et al*., 2016). Nevertheless, more than one in seven people globally still do not have access to sufficient protein and energy from their diets, with many suffering from some form of micronutrient deficiencies (FAO, 2009). This undesirable scenario is due to wide geographic variations in crop and livestock productivity, even across regions that experience similar climates (Godfray *et al*., 2010). In this connection, per capita food production in countries that experienced rapid economic growth, such as in Asia and Latin America, has increased approximately twofold over the past five decades. For instance, total meat production in the developing world tripled between 1980 and 2002, from 45 to 134 million tons (World Bank, 2009). In contrast, in Africa, per capita food production fell back from the mid-1970s and has only just reached the same level as in 1961 (Evenson and Gollin, 2003; FAOSTAT, 2009).

In 2017, The Kingdom of Eswatini (Swaziland)’s beef cattle population stood at 496 094 (DVLS, 2017), > 80% of which are owned by resource-poor smallholder farmers on Swazi National Land (SNL) (DVLS, 2014). The Nguni is the cattle breed raised by most livestock farmers on SNL (Nkondze *et al*., 2014) and comprises ∼ 70% of the cattle population (DVLS, 2004). There is also a sizeable number of exotic Brahman and Simmental breeds that have been introduced into the country for crossbreeding with the Nguni in order to improve beef production. Although the country has a beef quota to supply European markets, this quota has never been met due to the low off-take from the SNL. In order to boost beef production, small-scale farmers are being encouraged to venture into feedlotting of beef cattle and thus improve the contribution of the beef industry to the country’s economy (DVLS, 2004).

One major problem facing cattle producers, which therefore limits food production, in Eswatini and other tropical areas in Africa and elsewhere is poor nutrition of their animals especially during the dry season. Cattle grazing semi-arid range may be subjected to low intakes of metabolisable energy and protein as a result of drought, high stocking rates, or during the dry season low digestibility and low protein content of the forage (Butterworth, 1985). The growth and milk yield of cattle are reduced during such periods of under-nutrition (O’Donovan, 1984). It is during this season that problems such as disease and weight loss due to poor dietary regimes normally occur (Murphy and Colucci, 1999).

One way of improving livestock productivity in these areas involves the utilisation of locally available crop residues and by-products to formulate low-cost diets that are however adequate in essential nutrients such as energy and proteins (Kadzere, 1995; LDPS, 1998; Murphy & Colucci, 1999; Mlambo *et al*., 2011). In this regard, there are abundant quantities of agro-industrial by-products such as sugarcane tops, of which Eswatini produced approximately 920 000 metric tonnes in 2016/2017 (20% of sugarcane production, on wet basis) (Gain Report, 2016). Sugarcane tops can be utilized as livestock feed during this stressful period as they are highly palatable and rich in energy in the form of water-soluble carbohydrates. However, they are also deficient in and require supplementation with proteins and minerals (Abate *et al*., 1992; Mthiyane *et al*., 2001). Unfortunately dietary supplements formulated with grains and imported animal and plant-based conventional protein sources such as soyabean meal are too expensive and unaffordable by most smallholder farmers. With the feed cost of feedlotting animals being ∼ 70–80% of the total cost (Henning, 1999), these farmers are constrained from venturing into commercial feedlot production of beef. Hence the intense search for cheaper locally available yet nutritionally adequate alternative feed supplements.

Broiler litter is one of the most extensively used and intensively investigated cheap alternative protein (15% – 35% CP) and mineral rich supplementary feedstuffs for cattle and other livestock in Southern Africa (Mavimbela and van Ryssen, 2001; Mthiyane *et al*., 2001; Masaka *et al*., 2015). The utilisation of broiler litter is, however, limited by the presence of human pathogens (Chen and Jiang, 2014), pesticides and drug residues (Belewu, 1997) and heavy metals (Abdel-Mohsein and Mahmoud, 2015) which might pose health hazards to livestock and human consumers.

One safer and sustainable strategy of improving smallholder beef productivity in Africa and elsewhere involves dietary supplementation with the foliage of locally abundant and nutritious tropical leguminous and non-leguminous trees/shrubs. Tree/shrub foliage has been investigated as a potential supplement for ruminants because of its beneficial effect of increasing metabolizable energy intake, nitrogen intake, feed efficiency, animal performance (Osuji *et al*. 1995; Umunna *et al*. 1995; Teferedegne, 2000; Zhou *et al*., 2014; Adjorlolo *et al*., 2016; Hailemarium *et al*., 2016) and decreasing rumen fermentative production of methane gas that causes global warming (Gemeda and Hassen, 2015). Also, there is renewed interest in the use of tree foliage as a protein supplement in beef cattle production arising from beneficial manipulative effects of leaf phytochemicals on beef FA composition and enhancement of meat quality particularly in Southern Africa (Mapiye *et al*., 2011a, b & c; Qwele *et al*., 2013).

The increased interest in recent years in ways to manipulate the FA composition of beef arises from the growing consumer perception of the red meat as a major source of fat especially SFAs, which have been implicated in diseases associated with modern life. SFAs (bad), as opposed to their UFAs (good) counterparts, are generally associated with the worldwide proliferation of modern diseases of diabetes, cancer, cardiovascular disease and others (Scollan *et al*., 2001; Muchenje *et al*., 2009a; Vahmani *et al*., 2015). There is a growing body of evidence indicating that dietary supplementation with the foliage of various leguminous and non-leguminous trees alters the FA composition of ruminant meat away from the SFA towards the UFA (MUFA and PUFA) profile thus potentially enhancing the health status of consumers (Moyo *et al*., 2011; Marume *et al*., 2012; Qwele *et al*., 2013). This has led to interest in the determination of the total SFA, total MUFA, total PUFA as well as the MUFA/SFA, PUFA/SFA and PUFA/MUFA ratios of beef and other ruminant meats as indices of the health-related FA status of meat (Mapiye *et al*., 2011c; Marume *et al*., 2012). Also, the proportions, totals and ratios of the two classes of essential FAs (EFA), *n*-6 PUFA and *n*-3 PUFA, found in meat have been of interest as indices of the potential of meat to induce disease (Marume *et al*., 2012). High *n*-6 PUFA/*n*-3 PUFA ratios promote many chronic diseases whereas lower ratios have suppressive effects (Simopoulos, 2006; Vahmani *et al*., 2015). Further, quantification of the desaturase index that estimates the activity of steroyl-coenzyme-A-desaturase/Δ^9^-desaturase by comparing product-to-precursor FA ratios and is also important for production of CLA and MUFA (Souyert *et al*., 2006) has been used to compare and quantify SFAs and certain UFAs in ruminant products (Nantapo *et al*., 2014).

*M. azedarach* is a non-leguminous tree native to India, China and Iran but is now naturalized in many countries in Africa, Australia and America (Al-Rubae, 2009). It is often confused with *Azadirachta indica* (A.) Juss., another *Meliaceae* family plant species often called “neem”, due to their morphological similarity (Hördegen *et al*., 2003). *M. azedarach* is a hardy and drought resistant (Al-Rubae, 2009) as well as fast growing ornamental tree that grows to a height of 6–12 m, with some rainforest varieties reaching 30–45 m (Oelrichs *et al*., 1985; Hare, 1998). It has green leaves that turn yellow in the autumn and fall in the winter. The flowers are produced in large, sparse clusters in the spring and have a strong scent resembling the lilac scent (*Syringa vulgaris*). The fruits are cherry-like shape, yellowish to brown in colour and persist longer after the leaves have fallen (Stavarache *et al*., 2008).

*M. azedarach* is an invasive plant (Henderson, 1999, 2007) that also contains potentially toxic phytochemicals including azadirachtin (Mwandila, 2009), tetranortriterpene limonoids (Chung Huang *et al*., 1996; Carpinella *et al.*, 2003; Sanna *et al*., 2015), saponines (Oelrichs *et al*., 1985; Ahn *et al*. 1994; Nakatani *et al*. 1994; Huang *et al*. 1995), alkaloids (azadirine), resins, tannins and meliotanic and benzoic acids (Carratala, 1939). The toxic principles are mainly in the fruits, with their concentrations being lesser in the leaves (Hurst, 1942; Kingsbury, 1964; Everist, 1974; Kwatra *et al*., 1974). The intoxication by *M. azedarach* affects mainly pigs due to the ingestion of the fruits (Oelrichs *et al*., 1985; Timm and Riet-Correa, 1997) and the toxic dose for pigs is around 0.5% – 0.7% of the body weight (Hurst, 1942; Kingsbury, 1964). Cattle, sheep, goats and poultry can also be affected, but this is rare (Everist, 1974, Oelrichs *et al*., 1985; Hare, 1998). The mature fruits are more poisonous than the green ones (Hurst, 1942; Kingsbury, 1964). The toxicity of the plant could vary with the growing area and some trees can be non-toxic (Oelrichs *et al*., 1985). Indeed, no symptoms of toxicity were observed when goats were fed the plant’s foliage in India (Barman *et al*., 2003) and Tanzania (Damas Msaki *et al*., 2012), even when it was included in the caprine’s diet at as high a level as 60–70% in Vietnam (Hang *et al*., 2012). Also, oral administration of doses of the plant’s foliage or fruits induced no toxic effects in goats in Japan (Nakanishi *et al*., 2011) and Botswana (Madibela and Kelemogile, 2008), and in cattle in Bangladesh (Amin *et al*., 2008). Instead, benefits in terms of FI, BWG and decreased gastrointestinal nematode infestation were realized in these studies. Notwithstanding the toxicity, the plant has many good uses, primary among which is its medicinal value as an anthelmintic, tonic, anti-pyretic and also for the treatment of leprosy, eczema and asthma (Oelrichs *et al*. 1985). It is widely used as an anti-oxidative, analgesic, anti-inflammatory, insecticidal, rodenticidal, anti-diarrhoeal, de-obstruent, diuretic, anti-diabetic, cathartic, emetic, anti-rheumatic and anti-hypertensive (Sharma & Paul, 2013).

*M. azedarach* trees abundantly and widely occur in Southern Africa, including in Eswatini (Henderson, 1999, 2007). Its foliage is rich in protein (CP: 298.7 g/kg DM) and other nutrients (Ghazanfar *et al*., 2011; Gemeda and Hassen, 2015). Limited studies have shown positive effects of administration of MALM and its extract on body weight in cattle (Amin *et al*., 2008) and growth rate in goats (Datta *et al*., 2003). However, there are no studies that have investigated MALM as a potential supplementary feed for feedlot-fed beef cattle and, in particular, in relation to the health-related FA composition of beef. Therefore, the objectives of this study were to investigate the influence of dietary supplementation with MALM on productive performance, meat fat depot FA composition and health-related FA indices in Nguni x Brahman steers fed sugarcane tops based diets. The underlying hypothesis was that dietary supplementation with MALM would improve productive performance of steers and the profile of desirable FAs and health-related FA indices in their meat fat.

## Materials and Methods

### Location of study, source and preparation of materials

The study was conducted at the University of Swaziland (UNISWA) Luyengo campus farm located in the upper Middleveld of Swaziland at coordinates 26° 32′ south and 31° 14′ east, with an altitude of 600–800 m above sea level and a mean maximum and mean minimum temperatures of 23 °C and 11 °C, respectively. The annual rainfall ranges from 850 mm to 1000 mm (Monadjem and Garcelon, 2005), most of which occurs between October and April. Chemical analyses of MALM and experimental diets were performed in the Nutrition Laboratory of the Department of Animal Science, Luyengo Campus, UNISWA. Fatty acid and CP analyses were performed in the Lipid Chemistry Laboratory, University of Free State, South Africa.

*M. azedarach* leaves were harvested between October and December 2016 from naturally growing trees at Luyengo campus and around Malkerns. They were dried under shade with daily turning to avoid moulding in an open-sided shed with sufficient ventilation. They were then loaded onto a tractor-drawn trolley and transported for safe storage in a big shed with suffient ventilation. Also, immediately after harvesting, small (∼200 g) fresh samples of the leaves were fortnightly collected from the field and taken to the Nutrition Laboratory for drying (65 °C), further milling (1 mm) and chemical analysis.

Sugarcane tops were collected from a sugarcane plantation at Dalcrue Farm and dried for two weeks under sunlight with daily manual turning using a hand-held fork to facilitate drying. Hominy chop, mineral-vitamin premix and salt were supplied by Feedmaster (Pty) Ltd (Swaziland). Molasses was obtained from Royal Swaziland Sugar Corporation (RSSC) at Mhlume whilst broiler litter was from broiler houses outside Malkerns collected 4 days after the broiler batches were cleared. The broilers were of the Cobb 500 strain and had been reared in the houses for 6 weeks during which they were fed maize and soyabean based starter and finisher diets. The litter was sun-dried for almost 10 h/day for 4 consecutive days by spreading to a thickness of ∼ 1 cm on polythene sheets on a concrete floor. When the moisture content was around 10% – 15%, the litter was safely stored in 50 kg bags until it was incorporated into the experimental diets.

Dried *M. azedarach* leaves, sugarcane tops and broiler litter were then milled (13 mm sieve) using a tractor-mounted miller. Eighten (18) intact Nguni x Brahman crossbreed steers (18–20 months old; average initial liveweight = 250.0 ± 56 kg) were sourced from local smallholder farmers in KaBhudla, Ngogola, Thulwane and New Thulwane areas in the Lubombo region of Swaziland. Upon arrival at the University, steers were dewormed using Dectomax at the rate of 1 ml/10 kg body weight. They were also vaccinated against botulism using a 3 in 1 vaccine (Prondistar^®^). Dipping was done once before the trial using Paracide Dip to prevent tick-borne diseases. Further treatments were done when necessary.

### Experimental design, animals and diets

Eighteen (18) steers were used in a CRD experiment involving 3 dietary treatments with 6 replicate animals each. The experiment elapsed over 3 months, preceded by a 14-day adaptation period. The individual animal was the experimental unit. The animals were randomly assigned to experimental diets based on body weight. They were kept in individual stalls each with a pair of feeding and water troughs. Iso-energetic and iso-nitrogenous sugarcane tops and hominy chop-based diets, in meal form, formulated to meet the nutritional requirements of beef cattle as recommended by the NASEM (2016) and to which graded levels of MALM [0% (Control), 15% and 30%] were sequentially added in replacement of broiler litter (Table 1), were used.

**Table 1.** Ingredient and nutrient composition (%) of experimental diets

| Ingredient | *M. azedarach* leaf meal level |  |  |
| --- | --- | --- | --- |
|  | 0% | 15% | 30% |
| Sugarcane tops | 39.3 | 39.3 | 39.3 |
| Hominy chops | 20 | 20 | 20 |
| Broiler litter | 30 | 15 | 0 |
| *M. azedarach* leaf meal | 0 | 15 | 30 |
| Liquid molasses | 10 | 10 | 10 |
| Salt | 0.5 | 0.5 | 0.5 |
| Vitamin-mineral premix<sup>a</sup> | 0.2 | 0.2 | 0.2 |

<sup>a</sup>Mineral-vitamin premix supplied the following per kg of complete feed: vitamin A (retinol acetate) 4000 000 IU, copper 10g, manganese 30g, zinc 70g, iodine 0.8g, cobalt 0.3g and selenium 300 00mg.

### General management

Steers were kept in wooden pens (1 m × 2 m; 1 animal/pen) built inside a well-ventilated wooden mini-feedlot with sides open to natural light, raised wooden slab floor and a roof of corrugated iron sheets. Before commencement of the experiment, pens were disinfected with an antiseptic (Jeyes fluid). Steers were inspected daily for any health-related problems. Their diets were offered as a total mixed ration twice daily at 08h00 hrs and 17h00 hrs. General animal health was monitored by the UNISWA animal health specialists. The experimental diets were manually mixed weekly on a concrete floor, beginning with the smaller ingredients separately in molasses with some hominy chop and then combining these with the larger ingredients. Steers had *ad libitum* access to both feed and water, which were weighed and offered twice per day, in the morning at 7h00 and in the afternoon at 14h00. Feed refusals were recorded daily. The animals were weighed fortnightly in a crush pen using a 1000-kg scale that was first calibrated using standardised 50 kg weights.

In the evening of the last day of the experiment, all the steers were deprived of feed for 12 hrs before slaughter. Instead, they were provided with clean water only. The animals were slaughtered in the morning of day 91 at the UNISWA Farm abattoir. They were killed in a humanly manner as outlined by MoA (2012). After slaughter, bleeding and removal of hides, evisceration was done. About 1 kg samples of kidney fat were taken from each animal, placed in polystyrene bags and deep-frozen (−20 °C) before they were transported to Bloemfontein for analysis. All animals were cared for according to the animal care and welfare guidelines of MoA (2012).

### Measurements

Steers were individually weighed at the beginning (day 0) and then fortnightly thereafter until the end of the experiment. BWG was then calculated by subtracting the previous from the current weight and dividing by 14. Also, feed and water offered and left were weighed per pen daily in order to calculate average FI and WI per animal per day. FI and WI were therefore measured as the difference between feed or water offered and refusals. FCE was calculated as BWG/FI.

### Chemical assays

The DM (930.15), CP (954.01), ash (942.05), EE (920.39) and CF analyses of MALM and experimental diets were performed according to procedures of the AOAC (2000). CP was calculated using N × 6.25.

Total lipid from kidney fat was quantitatively extracted, according to the method of Folch *et al*. (1957), using chloroform and methanol in a ratio of 2:1. An antioxidant, butylated hydroxytoluene was added at a concentration of 0.001% to the chloroform: methanol mixture. A rotary evaporator was used to dry the fat extracts under vacuum and the extracts were dried overnight in a vacuum oven at 50°C, using phosphorus pentoxide as a moisture adsorbent. Total extractable intramuscular fat was determined gravimetrically from the extracted fat and expressed as % fat (w/w) per 100 g tissue. The extracted fat was stored in a polytop (glass vial, with push-in top) under a blanket of nitrogen and frozen at –20°C pending fatty acid analyses.

A lipid aliquot (±30 mg) of sausage batter lipid were converted to methyl esters by base-catalysed trans-esterification in order to avoid CLA isomerisation, with sodium methoxide (0.5 M solution in anhydrous methanol) during 2 h at 30 °C, as proposed by Park *et al*. (2001), Kramer *et al*. (2002) and Alfaia *et al*. (2007). Fatty acid methyl esters (FAMEs) from sausage batter lipid were quantified using a Varian 430 flame ionization GC, with a fused silica capillary column, Chrompack CPSIL 88 (100 m length, 0.25 mm ID, 0.2 μm film thicknesses). Analysis was performed using an initial isothermic period (40°C for 2 minutes). Thereafter, temperature was increased at a rate of 4°C/minute to 230 °C. Finally an isothermic period of 230 °C for 10 minutes followed. FAMEs n-hexane (1μl) was injected into the column using a Varian CP 8400 Autosampler. The injection port and detector were both maintained at 250 °C. Hydrogen, at 45 psi, functioned as the carrier gas, while nitrogen was employed as the makeup gas. Galaxy Chromatography Data System Software recorded the chromatograms.

FAME samples were identified by comparing the retention times of FAME peaks from samples with those of standards obtained from Supelco (Supelco 37 Component Fame Mix 47885-U, Sigma-Aldrich Aston Manor, Pretoria, South Africa). CLA standards were obtained from Matreya Inc. (Pleasant Gap, Unites States). These standards included: cis-9, trans-11 and trans-10, cis-12-18:2 isomers.

Fatty acids were expressed as the proportion of each individual fatty acid to the total of all fatty acids present in the sample. Fatty acid data were used to calculate the following ratios of FAs: total SFAs, total MUFAs, total PUFAs, PUFA/SFA, Δ^9^-desaturase index (C18:1*c*9/C18:0), total *n*-6 PUFA, total *n*-3 PUFA, the ratio of *n*-6 PUFA/*n*-3 PUFA. Atherogenicity index (AI) was calculated as: AI = (C12:0 + 4 x C14:0 + C16:0)/(MUFA + PUFA) (Chilliard *et al*., 2003).

### Statistical analyses

Data on productive performance, fatty acid composition, fatty acid totals and ratios and health indices were analysed using the GLM procedure of Minitab (2000). Statistical significance was accepted based on the 0.05 level of probability. Data are presented as least squares (LS) means with respective pooled standard errors of the mean (SEM). Where significant differences (p < 0.05) between treatments were observed, LS means were compared using the Tukey test.

## Results

The proximate composition of MALM is shown in Table 2. The tree foliage contained rather high CP (290.0 g/kg DM), CF (170 g/kg DM), EE (78.1 g/kg DM) and ash (77.5 g/kg DM) contents.

**Table 2.** The proximate composition of *M. azedarach* leaf meal

| Chemical component | Amount |
| --- | --- |
| DM (g/kg) | 291.0 |
| g/kg DM |  |
| CP | 290.0 |
| CF | 170.0 |
| EE | 78.1 |
| Ash | 77.5 |

The effects of dietary supplementation with MALM on productive performance and slaughter weights of beef steers are presented in Table 3. Both BWG and FCE were increased by dietary MALM supplementation (P < 0.01). In contrast, dietary MALM supplementation had no effect on FI, WI and slaughter weights of steers (P > 0.05). Also, there were no observable abnormal behaviours in the steers nor visible abnormalities on beef carcasses and internal organs.

**Table 3.** Body weight gain, feed and water intakes, feed conversion efficiency and slaughter weights of beef steers fed diets supplemented with graded levels of *M. azedarach* leaf meal

| Parameter | *M. azedarach* leaf meal level |  |  | Pooled SEM<sup>1</sup> | P-value |
| --- | --- | --- | --- | --- | --- |
|  | 0% | 15% | 30% |  |  |
| BWG (kg/d) | 0.77 | 2.62 | 3.27 | 0.20 | P < 0.01 |
| FI (kg/d) | 8.36 | 9.33 | 9.68 | 0.22 | P > 0.05 |
| WI (L/d) | 25.80 | 30.07 | 30.72 | 0.77 | P > 0.05 |
| FCE (BWG/FI) | 0.10 | 0.28 | 0.33 | 0.02 | P < 0.01 |
| Slaughter weight (kg) | 311.7 | 319.2 | 335.8 | 10.7 | P > 0.05 |

<sup>1</sup>SEM based on pooled estimate of variation (n = 6).

The influence of diet on the kidney fat depot SFA and UFA composition in beef steers is presented in Table 4. Overall, the predominant fatty acids in kidney fat of Nguni x Brahman steers were stearic acid (42.0%), palmitic acid (28.6%), oleic acid (18.1%), myristic acid (3.8%), linoleic acid (2.1%) and margaric acid (1.6%), in that order. Among SFAs, dietary MALM supplementation increased margaric acid (P < 0.05) whilst it decreased arachidic acid (P < 0.01) content. Otherwise, the diet had no effect on the content of all other SFAs (P > 0.05) in kidney fat. Regarding UFAs, dietary MALM supplementation increased kidney fat α-linolenic (P < 0.001) and CLA (P = 0.01) but had no effect on the content of all other UFA species (P > 0.05).

**Table 4.**
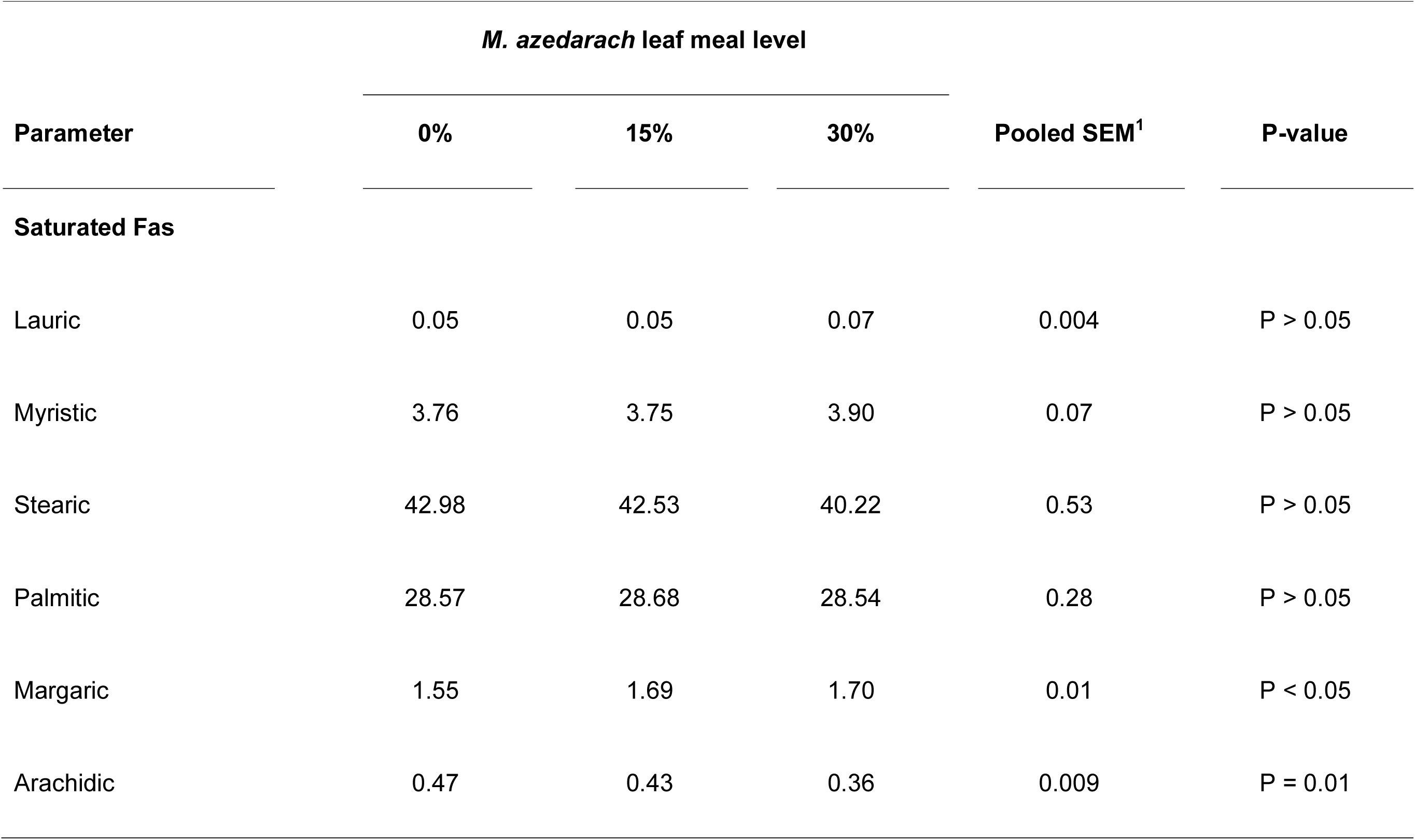
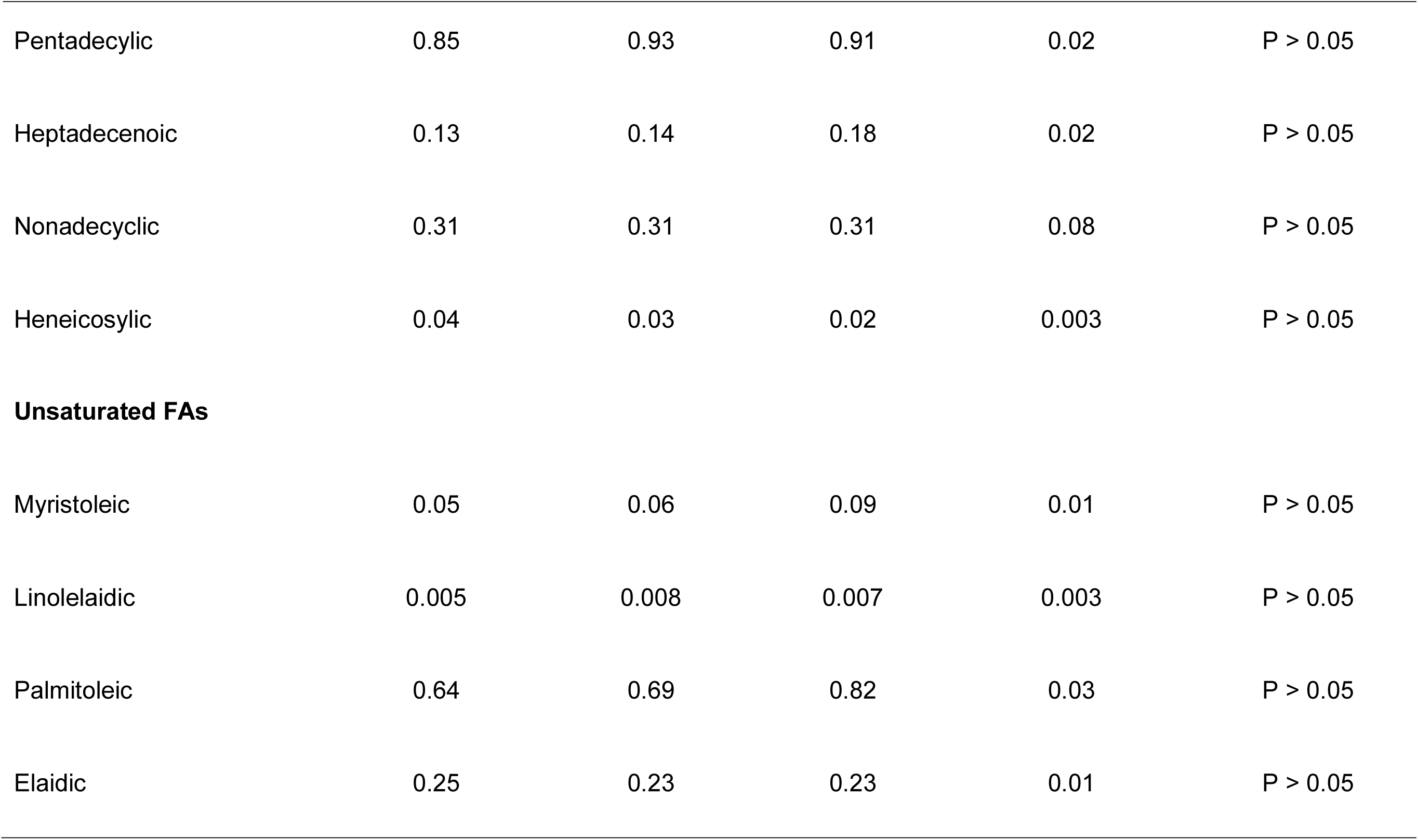
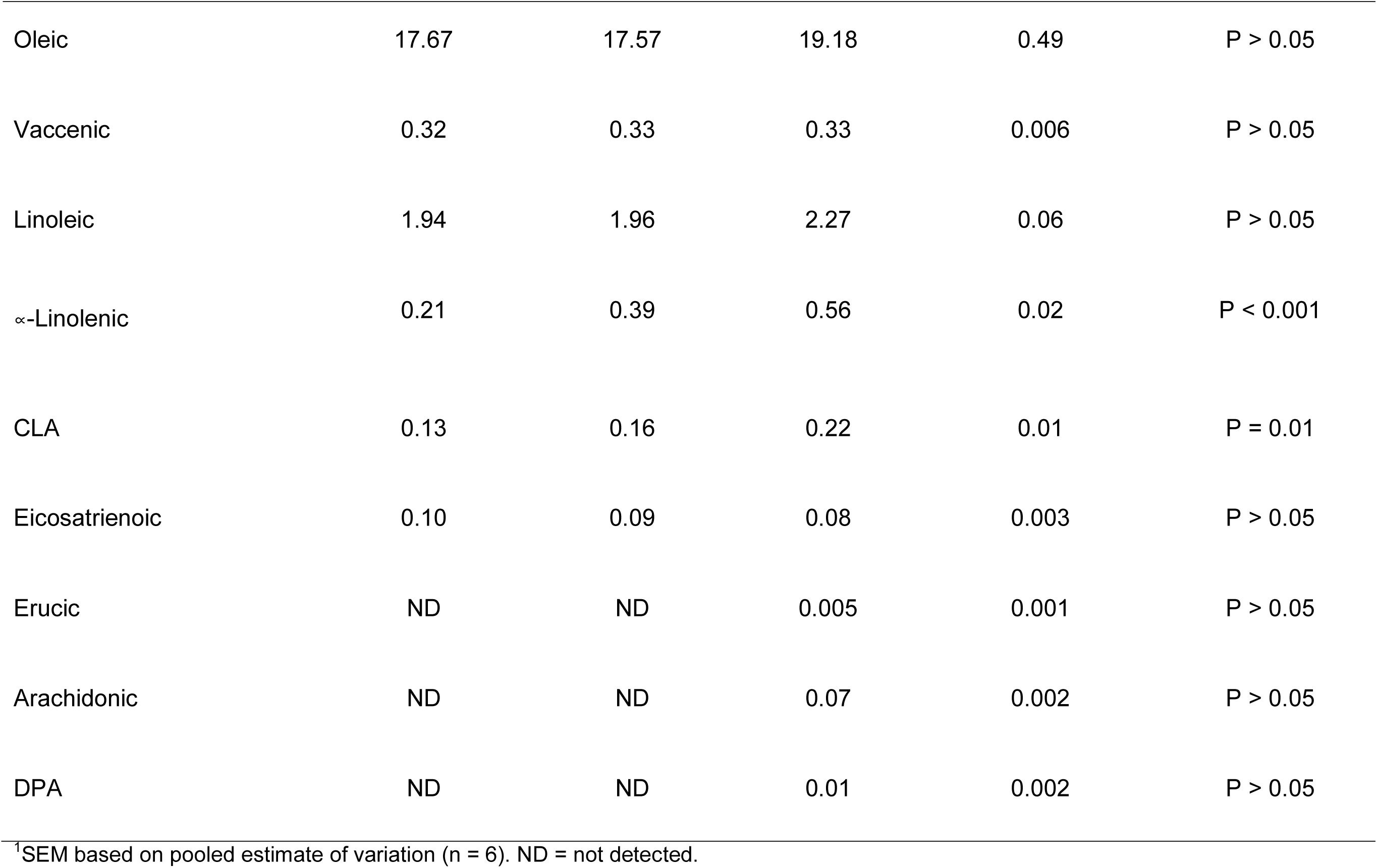
Kidney fat depot fatty acid composition in beef steers fed diets supplemented with graded levels of *M. azedarach* leaf meal

|  | *M. azedarach* leaf meal level |  |  |  |  |
| --- | --- | --- | --- | --- | --- |
| Parameter | 0% | 15% | 30% | Pooled SEM<sup>1</sup> | P-value |
| Saturated Fas |  |  |  |  |  |
| Lauric | 0.05 | 0.05 | 0.07 | 0.004 | P > 0.05 |
| Myristic | 3.76 | 3.75 | 3.90 | 0.07 | P > 0.05 |
| Stearic | 42.98 | 42.53 | 40.22 | 0.53 | P > 0.05 |
| Palmitic | 28.57 | 28.68 | 28.54 | 0.28 | P > 0.05 |
| Margaric | 1.55 | 1.69 | 1.70 | 0.01 | P < 0.05 |
| Arachidic | 0.47 | 0.43 | 0.36 | 0.009 | P = 0.01 |
| Pentadecylic | 0.85 | 0.93 | 0.91 | 0.02 | P > 0.05 |
| Heptadecenoic | 0.13 | 0.14 | 0.18 | 0.02 | P > 0.05 |
| Nonadecyclic | 0.31 | 0.31 | 0.31 | 0.08 | P > 0.05 |
| Heneicosylic | 0.04 | 0.03 | 0.02 | 0.003 | P > 0.05 |
| **Unsaturated FAs** |  |  |  |  |  |
| Myristoleic | 0.05 | 0.06 | 0.09 | 0.01 | P > 0.05 |
| Linolelaidic | 0.005 | 0.008 | 0.007 | 0.003 | P > 0.05 |
| Palmitoleic | 0.64 | 0.69 | 0.82 | 0.03 | P > 0.05 |
| Elaidic | 0.25 | 0.23 | 0.23 | 0.01 | P > 0.05 |
| Oleic | 17.67 | 17.57 | 19.18 | 0.49 | P > 0.05 |
| Vaccenic | 0.32 | 0.33 | 0.33 | 0.006 | P > 0.05 |
| Linoleic | 1.94 | 1.96 | 2.27 | 0.06 | P > 0.05 |
| α-Linolenic | 0.21 | 0.39 | 0.56 | 0.02 | P < 0.001 |
| CLA | 0.13 | 0.16 | 0.22 | 0.01 | P = 0.01 |
| Eicosatrienoic | 0.10 | 0.09 | 0.08 | 0.003 | P > 0.05 |
| Erucic | ND | ND | 0.005 | 0.001 | P > 0.05 |
| Arachidonic | ND | ND | 0.07 | 0.002 | P > 0.05 |
| DPA | ND | ND | 0.01 | 0.002 | P > 0.05 |

<sup>1</sup>SEM based on pooled estimate of variation (n = 6). ND = not detected.

The influence of diet on the kidney fat depot FA totals, health-related ratios and disease indices in beef steers is shown in Table 5. Dietary MALM supplementation increased kidney fat total PUFAs (P < 0.01), total *n*-3 PUFAs (P < 0.001), tended to increase the PUFA/SFA ratio (P = 0.08), decreased the *n*-6/*n*-3 PUFA ratio (P = 0.001) and increased the CLA/vaccenic acid index (P < 0.05). Otherwise, dietary MALM supplementation had no effect on all the other FA totals, ratios and health-related indices (P > 0.05).

**Table 5.**
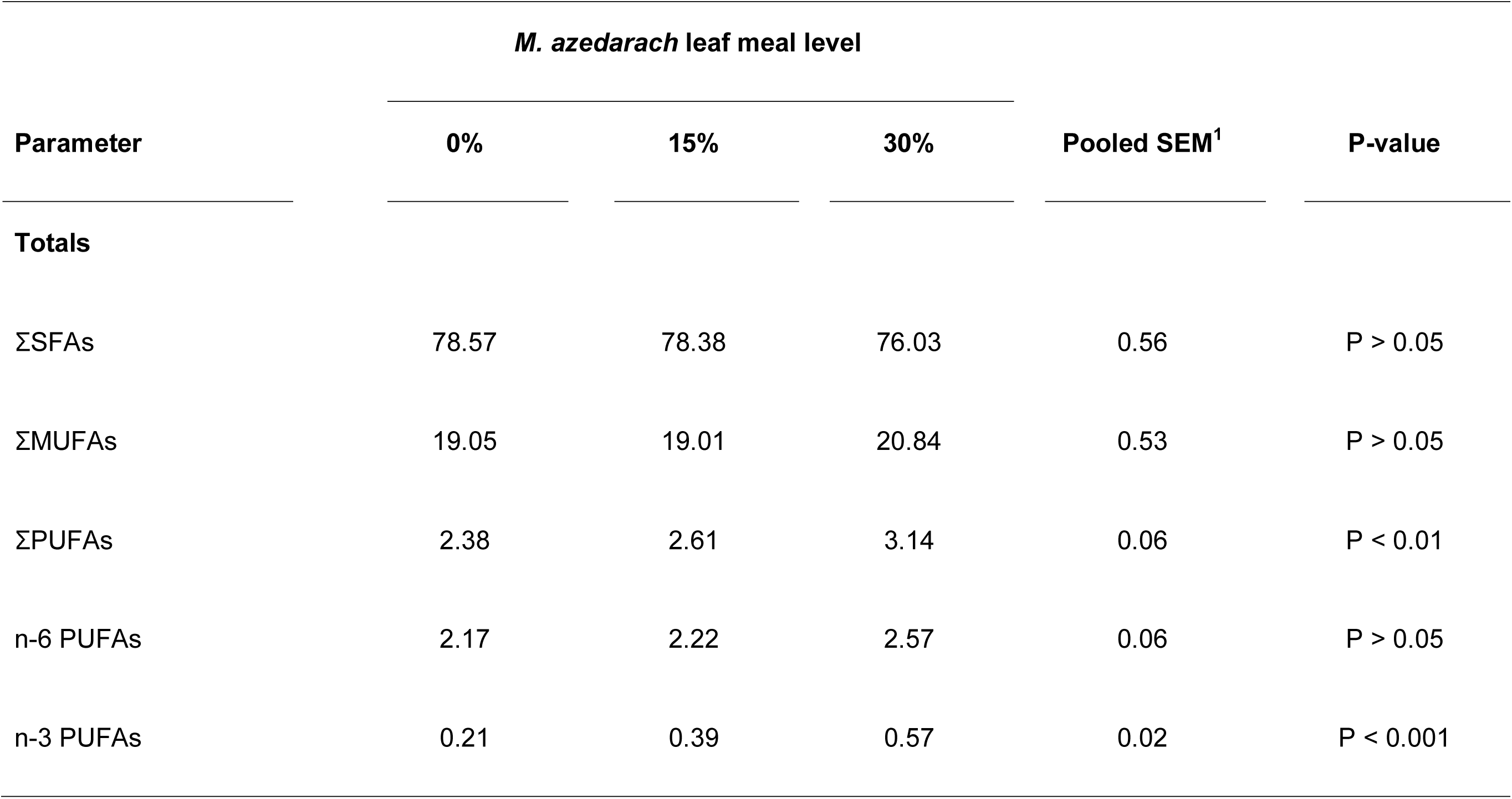
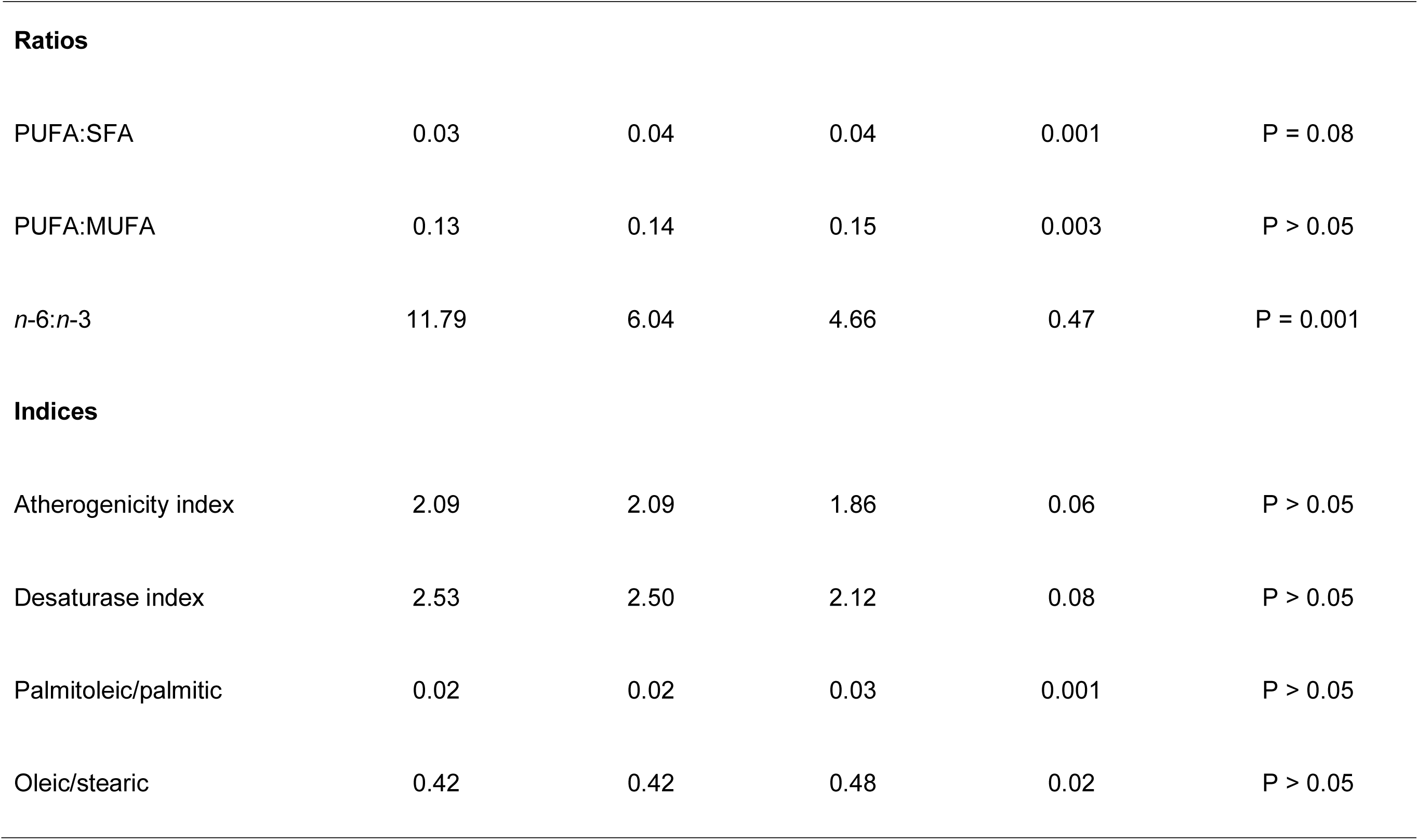
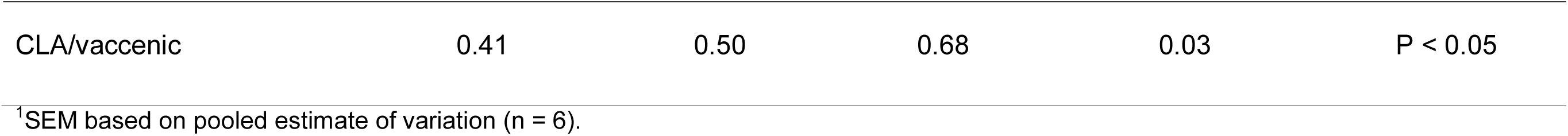
Kidney fat depot fatty acid totals and ratios and disease indices in beef steers fed diets supplemented with graded levels of M. azedarach leaf meal

| Parameter | *M. azedarach* leaf meal level |  |  | Pooled SEM<sup>1</sup> | P-value |
| --- | --- | --- | --- | --- | --- |
|  | 0% | 15% | 30% |  |  |
| **Totals** |  |  |  |  |  |
| ΣSFAs | 78.57 | 78.38 | 76.03 | 0.56 | P > 0.05 |
| ΣMUFAs | 19.05 | 19.01 | 20.84 | 0.53 | P > 0.05 |
| ΣPUFAs | 2.38 | 2.61 | 3.14 | 0.06 | P < 0.01 |
| n-6 PUFAs | 2.17 | 2.22 | 2.57 | 0.06 | P > 0.05 |
| n-3 PUFAs | 0.21 | 0.39 | 0.57 | 0.02 | P < 0.001 |
| **Ratios** |  |  |  |  |  |
| PUFA:SFA | 0.03 | 0.04 | 0.04 | 0.001 | P = 0.08 |
| PUFA:MUFA | 0.13 | 0.14 | 0.15 | 0.003 | P > 0.05 |
| *n*-6:*n*-3 | 11.79 | 6.04 | 4.66 | 0.47 | P = 0.001 |
| **Indices** |  |  |  |  |  |
| Atherogenicity index | 2.09 | 2.09 | 1.86 | 0.06 | P > 0.05 |
| Desaturase index | 2.53 | 2.50 | 2.12 | 0.08 | P > 0.05 |
| Palmitoleic/palmitic | 0.02 | 0.02 | 0.03 | 0.001 | P > 0.05 |
| Oleic/stearic | 0.42 | 0.42 | 0.48 | 0.02 | P > 0.05 |
| CLA/vaccenic | 0.41 | 0.50 | 0.68 | 0.03 | P < 0.05 |

<sup>1</sup>SEM based on pooled estimate of variation (n = 6).

## Discussion

Our results showed the foliage of *M. azedarach* to be very rich in CP, with relatively high contents of CF, EE and ash. Our results are in agreement with those (CP: 298.7 g/kg DM) of Gemeda and Hassen (2015), though these researchers found rather lower EE (30.1 g/kg DM) and slightly higher ash (112.7 g/kg DM) values. Azim *et al*. (2011) also found comparable CF (189.6 g/kg DM) but lower CP (14.1 g/kg DM), EE (40.4 g/kg DM) and ash (53.0 g/kg DM) values. In another study, higher ash (108–116 g/kg DM) but lower CP (184–251 g/kg DM) values have been found (Nassoro, 2014). The variations in nutritional content might be due to geo-climatic differences and stage of harvesting. Notwithstanding, MALM used in the current study contained very high CP content to qualify it as an alternative protein supplementary forage for beef cattle and other livestock. It is indeed a suitable substitute for broiler litter and other expensive conventional protein sources for beef cattle such as soyabean meal.

Our results also showed dietary MALM supplementation to have significantly increased BWG and FCE whilst it had no effects on FI, WI and slaughter weights in beef steers. This is the first study to demonstrate positive effects of dietary MALM supplementation in cattle. Hence, there are no comparative data in the literature. Notwithstanding, our results are in agreement with those of Damas Msaki *et al*. (2012) who observed significant increases in growth rate and BWG as well as milk production in goats (lactating does and growing kids) offered diets supplemented with 30% MALM for 3 months. Also, they concur with Hailemarium *et al*. (2016) who found supplementation with the foliage of neem, a closely related tree species, to significantly increase BWG, FCE, slaughter weight and other carcass characteristics in goats. Further, a water extract of *M. azedarach* foliage significantly increased body weight in cattle (Amin *et al*., 2008). Furthermore, Hang *et al*. (2012) found > 60% improvement in BWG in goats when the dietary *M. azedarach* content was raised from 60 to 70%. However, these researchers observed a decline in BWG when higher levels of the foliage were given, which indicated the presence of non-nutritional compounds. Indeed, *M. azedarach* toxicity has previously been observed in calves when its foliage was administered at doses ranging from 5–30 g/kg BW. The calves displayed clinical signs of depression, ruminal stasis, dry faeces with blood, ataxia, muscle tremors, sternal recumbency, hypothermia and abdominal pain, with some calves dying when dosed with 30 g/kg BW (del Carmen Méndez *et al*., 2002). No such untoward effects were observed even at 30% inclusion level of the foliage in the current study. The age of the animal might influence the sensitivity to toxicity, with calves likely to be more sensitive than adult cattle. In contrast to our data, Bakshi and Wadhwa (2007) found goat bucks to relish *M. azedarach* tree leaves as indicated by higher voluntary DM intake. These authors also found improved digestibility of nutrients as well as positive N-balance in the goats.

In terms of FA composition, our study showed the predominant FAs in kidney fat of Nguni x Brahman steers to be stearic acid (42.0%), palmitic acid (28.6%), oleic acid (18.1%), myristic acid (3.8%), linoleic acid (2.1%) and margaric acid (1.6%), in that order. This is in contrast to an observation whereby palmitic acid was the major FA followed by stearic acid and then oleic acid in Nguni steers (Mapiye *et al*., 2011c). This is perhaps not surprising as the Nguni x Brahman breed was used in our study, as opposed to the pure Nguni breed in the studies of Mapiye *et al*. (2011c). The Nguni x Brahman crossbreeds are now very common in Eswatini after Brahman bulls were introduced into the country some years ago in an attempt to improve beef production. The proportions of the major individual fatty acids were not within the normal range of values for beef cattle as reported by Muchenje *et al*. (2009b) and Alfaia *et al*. (2009), probably due to breed differences. However, the finding that stearic, palmitic and oleic acids were among the most abundant FAs in beef fat is consistent with previous reports (Partida *et al*., 2007; Muchenje *et al*., 2009b; Orellana *et al*., 2009).

Also, our results demonstrated that dietary MALM supplementation increased margaric acid whilst it decreased arachidic acid concentration of beef fat. This is the first study to demonstrate these effects of MALM on margaric and arachidic acids in beef fat. Hence, there are currently no comparable data in the literature. However, it would appear that MALM contained a high level of margaric acid for it to have so significantly influenced beef fat content of this SFA. In this regard, margaric acid has previously been found (at 0.4 x that of arachidic acid) in the seed oil of neem (Alaoui Ismaeli *et al*., 2016), a tree plant closely-related to *M. azedarach*. It is therefore possible that this SFA also occurs in *M. azedarach* foliage.

Both margaric and arachidic acids are SFAs. The former (also called heptadecanoic acid) is commonly present in bovine milk fat (Anderson, 1954; Mansson, 2008) and a variety of fish (Luzia *et al*., 2013) whilst the latter (as are behenic acid and ligonoceric acid) are contained in peanuts, canola oil, cashew nut and macadamia nuts (USDA-ARS, 2018). Whilst PUFA intake has been reported to be inversely associated with the risk of metabolic dysfunction (Shin *et al*., 2009; Kim *et al*., 2012), the intake of SFAs has been associated with adverse effects on the risk of metabolic syndrome (van Dijk *et al*., 2009; Xiao *et al*., 2006). In this regard, it is a well-established fact that palmitic acid (16:0) and stearic acid (18:0) have atherogenic and cardiovascular effects (French *et al*., 2002; Hodson *et al*., 2008). Also, there is evidence that odd-chain FAs such as margaric acid can build up in membrane lipids of nervous tissue, resulting in altered myelin integrity and demyelination, leading eventually to impaired nervous system functioning (Muller *et al*., 1998). However, recent studies have suggested that each SFA has different effects on metabolic conditions according to their chain length (Matsumoto *et al*., 2013). Thus, in contrast to entrenched beliefs that SFAs are bad for human health that are fostered by widespread recommendations for consumers to avoid high fat foods (including whole fat dairy products), higher erythrocyte membrane or plasma phospholipid margaric acid levels have been found to be potential protective factors against development of metabolic syndrome, type 2 diabetes and associated inflammation in humans (Krachler *et al*., 2008; Maruyama *et al*., 2008; Fernandes *et al*., 2013; Forouhi *et al*., 2014; Magnusdottir *et al*., 2014). Indeed, increased dietary intake of margaric acid decreases serum ferritin, the high level of which is associated with metabolic syndrome, and normalises blood glucose, triglycerides, and insulin (Venn-Watson *et al*., 2015). It is therefore desirable that dietary MALM supplementation increased beef fat margaric acid content. Also, consumption of SFAs with 20 or more C atoms - classified as very long chain SFAs (VLSFAs) – such as arachidic acid (20:0), behenic acid (22:0), and lignoceric acid (24:0) has been reported to have beneficial effects on the metabolic abnormalities, including diabetes and cardiovascular diseases (Matsumori *et al*., 2013; Lemaitre *et al*., 2014, 2015; Fretts *et al*., 2014; Forouhi *et al*., 2014). In this regard, Lee *et al*. (2015) recently demonstrated that higher intake of VLSFAs was significantly associated with favourable metabolic status, including lower levels of circulating triglyceride (TG). Also, they showed that higher intake of arachidic acid and total VLSFAs was negatively associated with metabolic syndrome risk.

Our results further demonstrated positive effects of dietary MALM supplementation on kidney fat α-linolenic and CLA contents. This is probably due to high content of these PUFAs in *M. azedarach* foliage. Indeed, Erdogan Orhan *et al*. (2012) found the leaf oil of *M*. *azedarach* to be dominated by linolenic (35.16%), palmitic (26.09) and linoleic (20.13%) acids, with lesser amounts of lauric (7.63%), myristic (4.07%), oleic (4.05%), stearic (1.86%) and caproic (1.02%) acids. This contrasted the FA profile of the fruit oil which was dominated by linoleic (62.76%) and oleic (24.42%) acids, with lesser amounts of palmitic (8.97%), stearic (2.84%) and caproic (1.02%) acids (Erdogan Orhan *et al*., 2012). Also, a previous study found the seed lipids of three plant species that belong to the *Meliaceae* family to contain the largest proportion (31-50%) of *cis*-vaccenic acid ever found in nature (Kleinman and Payne-Wahl, 1984). It is possible that *M. azedarach* foliage used in the current study also contained a high content of this PUFA. *Trans*-vaccenic acid (18:1*t*11), the predominant 18:1-*trans* isomer in grass-fed cattle (Dannenberger *et al*., 2004), is an intermediate produced during ruminal biohydrogenation of linolenic acid (18:3*n*-3) and linoleic acid (18:2*n*-6) in the process of biosynthesis of *cis*-9, *trans*-11 CLA (18:2*c*9*t*11) (Harfoot and Hazlewood, 1988). With *M*. *azedarach* foliage oil being rich in linolenic and linoleic acids (Erdogan Orhan *et al*., 2012), it is not surprising that dietary MALM supplementation increased beef fat CLA content in this study.

Both α-linolenic acid and CLA are very important *n*-3 PUFAs in relation to human health. α-Linolenic acid can be endogenously desaturated and elongated to long-chain *n*-3 PUFAs (*n*-3 LC-PUFA), *i.e.* EPA, DPA and DHA (Razminowicz *et al*., 2006) that have been demonstrated to exert various beneficial health effects including low rates of coronary heart disease, asthma, type 1 diabetes mellitus, multiple sclerosis, cancer, inflammatory bowel disease, rheumatoid arthritis and psoriasis (Simopoulos, 2008). Of the two physiologically important CLA isomers, rumenic acid (*cis*-9, *trans*-11 18:2 or 18:2*c*9*t*11) is the predominant isomer accounting for up to 80–90% of total CLA in ruminant products and is particularly believed to be beneficial for human health (Kramer *et al*., 1997; Razminowicz *et al*., 2006; Vatansever *et al*., 2000). In this regard, CLA has metabolic and health-promoting properties such as cancer prevention, decreased atherosclerosis, improved immune response and anti-diabetic properties (Whigham *et al*., 2000; Pariza, 2004; Palmquist *et al*., 2005). Also, CLA has antioxidant properties (Belury, 2002), decreases body fat, particularly abdominal fat, alters serum total lipids and decreases whole body glucose uptake (Thom *et al*., 2001). Therefore, the demonstrated positive effects of dietary MALM supplementation on beef fat α-linolenic acid and CLA content is of significant importance in terms of consumer health. Whilst previous studies indicated that pasture feeding of bulls on grass enhances the proportion of *n*-3 PUFAs and CLA of beef in comparison to feeding them on maize silage and concentrate (Dannenberger *et al*., 2004; Nürnberg *et al*., 2002), our results suggest that beef animals can still be reared intensively but supplemented with the foliage of *M. azedarach* through their diets in order to enhances their meat content of *n*-3 PUFAs and CLA.

The positive effects on beef fat α-linolenic and CLA contents concur with the observed increase in the CLA/vaccenic acid index following dietary MALM supplementation in this study. Unfortunately, no comparative data exists in the literature as this is the first study to show the influence of tree foliage based diets on the CLA/vaccenic acid ratio in ruminants. Notwithstanding, our results corroborate a previous *in vitro* study whereby leaf extracts of *A. indica*, another *Meliaceae* tree plant closely related to *M. azedarach*, enhanced ruminal fluid content of rumenic acid (*cis*-9, *trans*-11 CLA), vaccenic acid and the ratio of CLA plus vaccenic acid to stearic acid (Heidarian Miri *et al*., 2013). It is therefore evident that both *A. indica* and *M. azedarach* have immense capacity to favourably influence ruminal biohydrogenation processes resulting in enhanced production of CLA, probably through their high content of linoleic and α-linoleic acids (Erdogan Orhan *et al*., 2012; Rao *et al*., 2014; Akacha *et al*., 2016). Indeed, CLA isomers are formed endogenously by the action of Δ^9^-desaturase on vaccenic acid (*trans11* C18:1), which is another product of biohydrogenation of linoleic acid and α-linoleic acid in the rumen (Harfoot and Hazelwood, 1997; Griinari *et al*., 2000).

Furthermore, our data demonstrated dietary *M. azedarach* foliage induced improvements in beef fat total PUFAs, total *n*-3 PUFAs and PUFA/SFA ratio as well as a decrease in the *n*-6/*n*-3 ratio. Again, this is the first study to present evidence of beneficial effects of dietary MALM supplementation on human health-related FA totals and ratios. In a previous study by Mapiye *et al*. (2011c), similar significant improvements in PUFAs and total *n*-3 PUFAs proportions were observed in meat from steers fed a diet supplemented with *Acacia karroo* leaf meal. These researchers found meat from steers fed a diet supplemented with *Acacia karroo* leaf meal to have higher PUFA/MUFA and PUFA/SFA ratios than meat from steers fed a sunflower cake supplemented diet. Also, the lowest *n*-6/*n*-3 PUFA ratio was recorded in meat from steers that received the *Acacia karroo* diet (Mapiye *et al*., 2011c). Further, Qwele *et al*. (2013) observed improved total PUFA, total *n*-3 PUFAs, PUFA/SFA ratio and a decrease in the *n*-6/*n*-3 PUFA ratio in the meat from goats fed *Moringa oleifera* leaf meal-supplemented diets. The *n*-6/*n*-3 PUFA ratio plays an important role in reducing the risk of coronary heart disease (American Heart Association, 2008). Our data showed beef fat from steers fed the 30% MALM supplemented diet to have a *n*-6/*n*-3 PUFA ratio of 4.66:1 which is below the recommended maximum of 5:1 (Razminowicz *et al*., 2006). In contrast, the meat fat from steers whose diets were supplemented with broiler litter had a *n*-6/*n*-3 PUFA ratio of 11.79:1, which is way above the recommended maximum and is a clear indication that feeding broiler litter as a protein supplement to beef cattle is not good in terms of meat quality and consumer health.

## Conclusion

This study showed *M. azedarach* foliage to be a potential alternative protein-rich feedstuff for beef cattle. Dietary supplementation with the foliage improved productive performance without influencing feed and water intakes. Also, by increasing the beef fat content of health-beneficial very long chain SFAs, the essential FA α-linolenic acid, CLA, total PUFAs, total *n*-3 PUFAs and CLA/vaccenic acid index whilst decreasing the *n*-6/*n*-3 PUFA ratio, it enhanced the eating quality of beef and demonstrated potential to enhance the health status of consumers. *M. azedarach* foliage therefore shows great potential for utilization as a cheap alternative protein-rich supplementary feedstuff that can replace broiler litter and other expensive conventional protein sources for beef cattle.

## Acknowledgements

We are grateful to the International Livestock Research Institute (ILRI) for financial support to undertake this study, UNISWA Transport Department for providing transport to ferry sugarcane tops and all feed ingredients from suppliers, Dalcrue Farm for the supply of sugarcane tops, Royal Swaziland Sugar Corporation for supplying molasses, Feedmaster (Pty) Ltd for supplying hominy chop, vitamin-mineral premix and salt, Mr. Mshumbu Mkhabela for helping with animal slaughtering, and Mr. Kuhle Ntshalintshali for helping with mixing of experimental diets, feeding and management of animals.

## Compliance with ethical standards

The care and use of steers were performed following the ethical guidelines of the UNISWA Department of Animal Science Board that approved the protocol used in the experiment.

